# Auditory Stimulus-response Modeling with a Match-Mismatch Task

**DOI:** 10.1101/2020.11.05.370072

**Authors:** Alain de Cheveigné, Malcolm Slaney, Søren A. Fuglsang, Jens Hjortkjaer

## Abstract

An auditory stimulus can be related to the brain response that it evokes by a stimulus-response model fit to the data. This offers insight into perceptual processes within the brain and is also of potential use for devices such as Brain Computer Interfaces (BCI). The quality of the model can be quantified by measuring the fit with a regression problem, or by applying it to a classification task and measuring its performance. Here we focus on a *match-mismatch* (MM) task that entails deciding whether a segment of brain signal matches, via a model, the auditory stimulus that evoked it. The MM task allows stimulus-response models to be evaluated in the limit of very high model accuracy, making it an attractive alternative to the more commonly used task of auditory attention detection (AAD). The MM task does not require class labels, so it is immune to mislabeling, and it is applicable to data recorded in listening scenarios with only one sound source, thus it is cheap to obtain large quantities of training and testing data. Performance metrics from this task, associated with regression accuracy, provide complementary insights into the relation between stimulus and response, as well as information about discriminatory power directly applicable to BCI applications. Using these metrics, we describe a range of models of increasing complexity that we compare to methods in the literature, showing state-of-the-art performance. We document in detail one particular implementation, calibrated on a publicly-available database, that can serve as a robust reference to evaluate future developments.

## Introduction

Continuous stimuli such as speech or music elicit an ongoing brain response (Ahissar et al., 2001; Aiken and Picton, 2008; Power et al., 2011; Ding and Simon, 2012; Kubanek et al., 2013) that can be detected with electroencephalography (EEG) or magnetoencephalography (MEG). The relation between stimulus and response can be characterized by fitting a model to the data (Lalor et al., 2009; Crosse et al., 2016). Most work has used a *linear* stimulus-response model to relate some feature transform of the stimulus (envelope, spectrogram, etc.) to the brain response. Such models come in three main flavors: a *forward model* that attempts to predict the neural response from the stimulus (Lalor et al., 2009; Ding and Simon, 2012; Crosse et al., 2016), a *backward model* that attempts to infer the stimulus from the response (Mesgarani and Chang, 2012; O’Sullivan et al., 2015; Puvvada and Simon, 2017; Hausfeld et al., 2018; O’Sullivan et al., 2019; Akbari et al., 2019), or a *hybrid forward-backward model* that transforms both stimulus and response to better reveal their relation (Dmochowski et al., 2017; de Cheveigné et al., 2018; Zhuang et al., 2020). The fit of the model is usually quantified by calculating the correlation coefficient between stimulus and response: the observation of a significant correlation suggests that the model captures some aspect of neural processing, and details of the model (e.g. latency or shape of a temporal response function) then provide insights into the sensory processing mechanisms at work within the brain.

The stimulus-response model can also be applied to a classification task, and its quality evaluated based on performance in that task. Auditory attention decoding (AAD) has played an important role in past studies (Kerlin et al., 2010; Power et al., 2011; Ding and Simon, 2012; Mesgarani and Chang, 2012). A subject is instructed to attend to one of two concurrent streams, usually speech, and the algorithm decides which stream was attended based on the brain activity (Fig. 1, left). Model accuracy can be quantified by classification performance (for example percent correct), and the task itself may be relevant for applications such as controlling a device (for example a hearing aid) based on which stream is the focus of attention. However, AAD requires a two-voice stimulus, specific instructions to subjects, and a well-controlled experimental setup. Data for training and evaluation depend on labels defined by the experimental task (specifically which voice the subject is attending). The listener’s attentional state may stray momentarily from instructions (e.g. attentional capture by the “unattended” stream) and so some proportion of the data may be mislabeled. This can be a problem if we wish to evaluate algorithms in the limit of small error rates.

**Figure 1:**
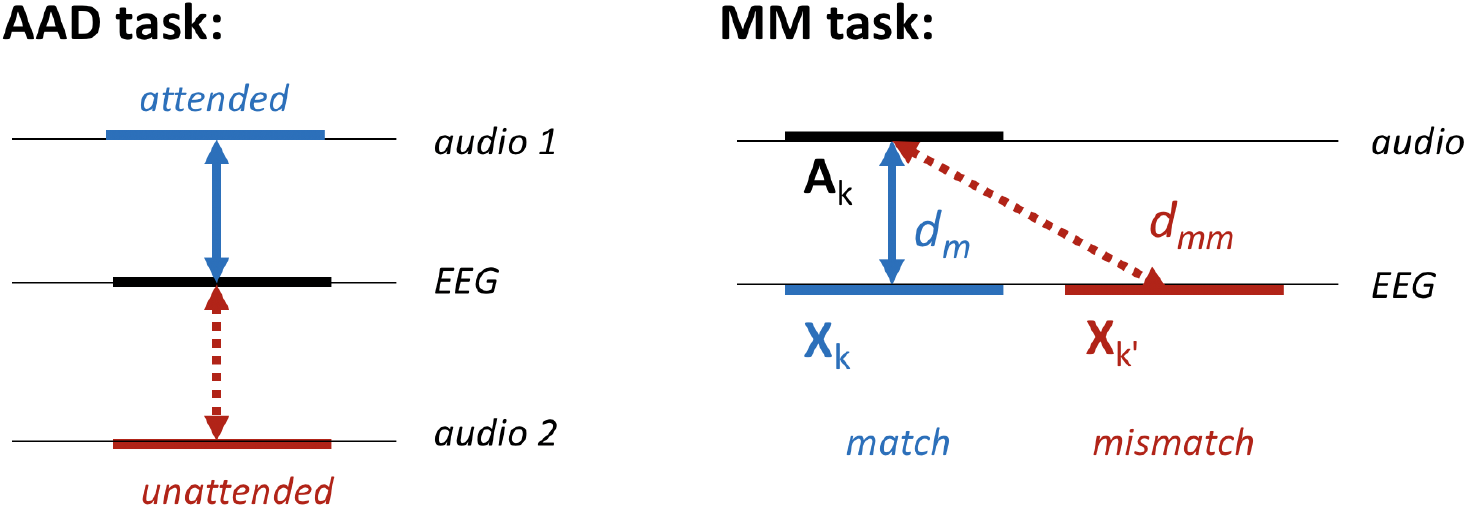
Left: the classic auditory attention detection task (AAD). A segment of EEG data is compared, via a model, to two audio streams, one attended and the other unattended. Right: the new match-mismatch task (MM). A segment of audio is compared, via a model, to the segment of EEG that it evoked (match, Euclidean distance *d*_*m*_) or some unrelated segment (mismatch, *d*_*mm*_).

In this paper, we consider a different classification task (*match-mismatch*, MM) that applies to listening scenarios with only one sound source. This task consists of deciding whether a segment of EEG or MEG is temporally aligned with a segment of audio (i.e. that segment of response was evoked by that paricular segment of stimulus), or not (Fig. 1, right). Compared to AAD, the MM task offers a simpler and potentially more efficient framework to optimize stimulus-response models. It can be applied in listening scenarios where there is only one speaker, and does not depend on whether the listener followed instructions as to which stream to attend (variations in attention to the single stream are possible but less disruptive). As no data labels are required, models may be trained for this task in a self-supervised manner. This simpler task is applicable to the evaluation of high performance algorithms with small error rates. To the extent that both MM and AAD rely on the accuracy of the stimulus-response models, we speculate that models optimized with one may yield improved performance on the other. Using MM, performance can be quantified by either the *sensitivity index*, defined here as the standardized mean of the distribution of the decision metric, or the *error rate*. Together, correlation, sensitivity index, and error rate form a trio of complementary performance metrics of stimulus-response models.

Building on prior work, cited above, we introduce a set of refinements applicable to a stimulus-response model and evaluate them within the MM task framework. These refinements allow more complex models while controlling for over-fitting. As we will show, error rates averaged over subjects for 5 s segments fall from ∼30% for our simplest model to ∼3% for the best (0% error for a subset of subjects) indicating considerably more reliable stimulus-response models. We devote effort to understanding which processing steps improve performance, and why. In the past, progress has been slowed by the lack of reliable comparative evaluation due to the diversity of experimental conditions and data, the absence of state-of-the-art algorithms in the “line-up”, and the aforementioned issues with associated with the AAD task. We use a publicly available database, metrics based on the simpler MM task, and a well-defined implementation of a competitive method to facilitate evaluation of future advances.

This study offers two main contributions. First, it introduces a simple objective task, match-mismatch (MM), to help in the evaluation of stimulus-response models. Second, it documents a set of techniques that boost performance beyond state of the art.

## 1 Methods

This section describes the stimulus-response model and provides details of the evaluation methods and experiments. The busy reader is encouraged to read the Section 1.1, then skip to Section 2, Results, and come back for more details as needed. We assume that brain responses are recorded by EEG, but the same methods are applicable to MEG or other recording modalities.

### 1.1 Models and metrics

In this subsection we define the mathematical tools to describe what we wish to accomplish, and the metrics to judge success.

#### Data Model

The brain response data consist of a time series matrix **X** of dimensions *T* (time) × *J* (channels). Each channel is the weighted sum of brain sources of interest as well as undesired noise and artifacts:

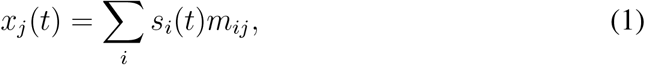

where *t* is time, [*s*_*i*_(*t*)], *i* = 1 *… I* are sources, and the *m*_*ij*_ are unknown source-to-sensor mixing weights. In matrix notation **X**=**SM**. This matches the physical source-to-sensor mixing process which is, to a good approximation, linear and instantaneous. The audio stimulus is represented as a matrix or column vector **A**, usually a transform such as the waveform envelope (akin to a measure of “instantaneous loudness”) or the spectrogram (akin to an “auditory nerve activity pattern”). **A** is of size *T* × *K*, where *K* is the number of channels of the stimulus representation (e.g. number of frequency bands of a spectrogram). In the following, *K* = 1.

#### Stimulus-Response Model

We assume that a transform **F** of the stimulus representation is non-trivially related to a transform **G** of the EEG:

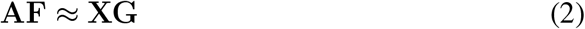

where ≈ indicates similarity according to some metric. By non-trivial we mean that Eq. 2 can be used empirically to decide whether or not a segment **X**_*s*_ of the brain data was recorded in response to a segment **A**_**s**_ of the stimulus. **F** and **G** are linear transform matrices (possibly convolutive), but Eq. 2 can be usefully generalized to more complex transforms, e.g. *f*_*X*_(**A**) ≈ *g*_*A*_(**X**).

Three special cases are worth noting. In the *forward model*, **AF** ≈ **X**, the transform **F** is used to predict the response from the stimulus. In the *backward model*, **A** ≈ **XG**, the transform **G** is used to infer the stimulus from the response. Forward and backward models are also referred to as “encoding” and “decoding” (Naselaris et al., 2011), or “temporal response function” (TRF) and “stimulus reconstruction” models, respectively. A third *hybrid* model involves transforms of both: **AF** ≈ **XG**. Tradeoffs between these three approaches are reviewed in the Discussion.

The transforms **F** and/or **G** are found by a data-driven algorithm, regression for the first and second models, or canonical correlation analysis (CCA) for the third. Given datasets **A** and **X**, CCA finds transforms such that (a) columns of **AF** are orthonormal (variance 1 and mutually uncorrelated), (b) columns of **XG** are orthonormal, (c) the pair formed by the first column of **AF** and the first column of **XG** has the greatest possible correlation on the training data, the pair formed by the second columns has the greatest correlation once the first columns have been projected out, and so-on. Matrices **F** and **G** are of size *J* × *H* and *K* × *H* respectively, where *H* is at most equal to the smaller of *J* and *K*.

#### The Match-mismatch Task

To assist evaluation, we define the match-mismatch (MM) task as follows. Given a segment of stimulus signal **A**_**s**_, the segment of EEG signal **X**_**s**_ that it evoked, and some unrelated segment of EEG signal **X**_**s**_*′ ≠***s**, decide which of the two EEG segments matches, via a model, the stimulus (Fig. 1, right). A practical application might be to determine whether a user is attentive to sound, or whether a particular alarm sound was noticed. Here we use it simply to mesure the accuracy of the stimulus-response model.

#### Metrics

Goodness-of-fit will be evaluated using three metrics: correlation, sensitivity index, and classification error rate, the last two contingent on the MM task. The first, correlation, is calculated between transforms **AF** and **XG** over the full duration of the data, or over a shorter segment of duration *D*. When the transformed data are normalized, as they are in this paper, correlation is related to Euclidean distance by the relation *r* = 1 − *d*^2^*/*2. A perfect match is characterized by *r* = 1, *d* = 0 and lack of correlation by (in expectation) *r* = 0, 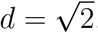.

The second metric, sensitivity index, is based on the distribution of the difference Δ_*s*_ = *d*_*mm*_ − *d*_*m*_ of Euclidean distances for matched and mismatched segments. For each segment *s* of stimulus (transformed and z-scored), *d*_*mm*_ is calculated as the average distance to mismatched segments *s*^*′*^ of EEG (transformed and z-scored), over all *s*^*′*^ *≠ s*, while *d*_*m*_ is the distance to the matched segment of EEG features. Values of *d*_*mm*_ cluster around 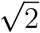 because the data are normalized and mismatched segments are uncorrelated. Matched distances *d*_*m*_ tend to have smaller values, and so the difference Δ_*s*_ is (hopefully) positively distributed. The sensitivity index is calculated as the mean of this distribution divided by its standard deviation (standardized mean):

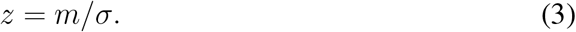

This definition is analogous to that of the “standardized mean difference” (d-prime), but differs in that it quantifies the *distribution of the difference* between *d*_*m*_ and *d*_*mm*_, rather than the distributions of those values themselves. For CCA, the distances are calculated taking into account the first 5 CCs with equal weight. In a previous study (de Cheveigné et al., 2018) we used linear discriminant analysis to optimally combine CCs, here we found less benefit so we report only this simpler scheme.

The third metric, error rate, counts the proportion of segments classified incorrectly in the MM task (the proportion of segments *s* for which Δ_*s*_ *<* 0). Sensitivity and error rate depend on segment duration *D*, which is varied as a parameter: the shorter the segment, the noisier the correlation or decision calculation, and the harder the task. Error rate (*e*) is preferred to proportion correct (1 − *e*) because, plotted on a logarithmic scale, it better reveals incremental steps towards better performance, particularly in the region of high performance. Each metric has its virtues, as elaborated in Section 3, Discussion.

#### Cross-validation

Regression and CCA are data-driven and thus prone to over-fitting. To avoid inflating results, a model can be trained and tested on different sets of data, for example using 𝒦-fold cross-validation (Murphy, 2021). Data are divided into 𝒦 trials, the model is fit on 𝒦 − 1 trials and performance metrics are evaluated on the *k*th (left-one-out), the final score being the average of these 𝒦 estimates.

### 1.2 Extending and reducing the model

At least three factors degrade the model fit: *latency, spectral mismatch* between the stimulus representation and the brain response, and *additive noise* in the response. These can be addressed in part by augmenting the data with a set of time lags (or a filter bank).

#### Lags and Time Shift

Brain responses unfold over time, and there may be a convolutive mismatch in the process connecting stimulus and response. These can be absorbed by augmenting the stimulus and/or brain signals with *time lags*. Applying a set of lags 0 *… L*_*A*_ − 1 to **A** and concatenating the time-lagged channels side by side yields a matrix of size *T* × *KL*_*A*_. Similarly, applying *L*_*X*_ lags to **X** yields a time-lagged matrix of size *T* × *JL*_*X*_. An appeal of lags is that they allow the algorithm (univariate regression or CCA) to automatically synthesize a *finite impulse response filter* (FIR) or, in the case of multichannel data, a multichannel FIR. This allows the model to minimize spectral mismatch (amplitude and phase) between **A** and **X**. The number of lags *L* determines the order of the synthesized FIR filter. A larger *L* confers the ability to select or reject temporal patterns on a longer time scale (lower frequencies), at the cost of greater computational cost and greater risk of overfitting.

In addition to lags, we introduce an overall *time shift S* between stimulus and response. This parameter, distinct from the lags, is intended to absorb any gross temporal mismatch due to instrumental or sensory latencies. This frees the lag parameters to fit finer spectro-temporal characteristics. Without *S* a larger value of *L* might be needed, with greater risk of overfitting. *S* is treated as a hyperparameter: the fit is repeated for several values and the one that yields the highest correlation value is retained.

#### Dyadic filter basis

Lags 0 *… L* − 1 form a basis of the space of FIR filters of order *L*, but one can choose a different basis, for example outputs of a *L*-channel *filter bank* of FIRs of order *L*. To reduce dimensionality, one can then choose a subset *L*^*′*^ *< L* of that basis, defining a *L*^*′*^-dimensional *subspace* of the space of FIRs of order *L*. With a judicious choice of filter bank, performance with *L*^*′*^ *< L* channels may be superior to merely choosing *L*^*′*^ *< L* lags, in part due to a lower risk of overfitting. For example, a logarithmic filter bank (e.g. wavelet, or dyadic) can capture patterns of both short and long time scale with a limited number of channels, whereas capturing the same long time scale with a basis of lags would entail a much larger dimensionality. Here, we use a dyadic filter basis.

#### Dimensionality reduction

The models we describe here can be large, including a large number of parameters, leading to overfitting if we do not have enough training data. Overfitting can be made less severe by *reducing the dimensionality* of the data before fitting the model, or by applying *regularization* within the fitting algorithm (Wong et al., 2018). The two approaches are closely related (Tibshirani et al., 2017, Sect 3.4.1). Here, we use dimensionality reduction. Data are submitted to Principal Component Analysis (PCA) and principal component (PCs) beyond a certain rank *N* are discarded, thus ignoring directions of low variance within the data. Ridge regularization similarly shrinks low-variance directions (Tibshirani et al., 2017).

### 1.3 Evaluation

Given the task described above, there are several ways we can measure success. This subsection describes the methodology, using cross-validation to limit overly optimistic measures of success.

#### Data

The data we use here are from a study that aimed to characterize cortical responses to speech for both normal-hearing and hearing-impaired listeners (Fuglsang et al., 2020). Experimental details are provided in that paper, and the data themselves are available from http://doi.org/10.5281/zenodo.3618205. In brief, 64-channel EEG responses to acoustic stimuli (audiobook) were recorded at a sampling rate of 512 Hz from 44 subjects, including both normal-and hearing-impaired. Stimuli for the latter were equalized (frequency-specific amplitude boost) to compensate for the impairment, and we pool data from both. Including both populations results in a larger and more diverse data set, with results possibly valid for a wider population (some applications target impaired users). Stimuli presented to each subject included 16 segments of single-talker speech with a male or female talker speaking in quiet, each of 50 s duration, that we consider in this study. Other stimuli presented in the same recording session (concurrent speech, tones) are not used. The publicly available dataset includes the temporal envelope of the speech stimulus, sampled at the same rate as the EEG, calculated by a model of instantaneous loudness that has been shown to be a predictor of cortical responses (Lalor et al., 2009; Ding and Simon, 2012; Di Liberto et al., 2015; Crosse et al., 2016).

#### Preprocessing

The EEG data were smoothed by convolution with a square window of duration 1/50 Hz (implemented with interpolation) to suppress the line artifact (50 Hz and harmonics) and downsampled by smoothing with a 4-sample square window and decimation by a factor of 4 to 128 Hz. The data were de-trended by applying a robust detrending algorithm (de Cheveigné and Arzounian, 2018) that robustly fit a 2nd order polynomial to overlapping intervals of size 15 s, subtracted the fit, and “stitched” detrended intervals together with a standard overlap-add procedure. The data were then high-pass filtered at 0.5 Hz using an order-2 Butterworth filter, then low-pass filtered at 30 Hz also with an order-2 Butterworth filter, and cut into 16 trials of 50 s duration. To remove eyeblink artifacts, a temporal mask was derived from the absolute power on a combination of two EOG channels and three frontal channels (F1, F2, Fz). Using this mask as a bias, the DSS algorithm was applied to find a transform maximizing eyblink activity (de Cheveigné and Parra, 2014) and the first two components (representing eyeblink artifact) were projected out of the EEG data.

To avoid aggravating the mismatch between stimulus and brain response, the stimulus envelope was filtered using the same high pass and low pass filters as for the EEG. All filters were “single pass” (causal).

#### Basic Models

To ease comparison with other studies, we define six models (Fig. 2) that illustrate basic processing choices, some of which have been made in prior studies and all of which are useful to understand in detail. For each, an overall time shift *S* is applied to the stimulus relative to the EEG.

**Figure 2:**
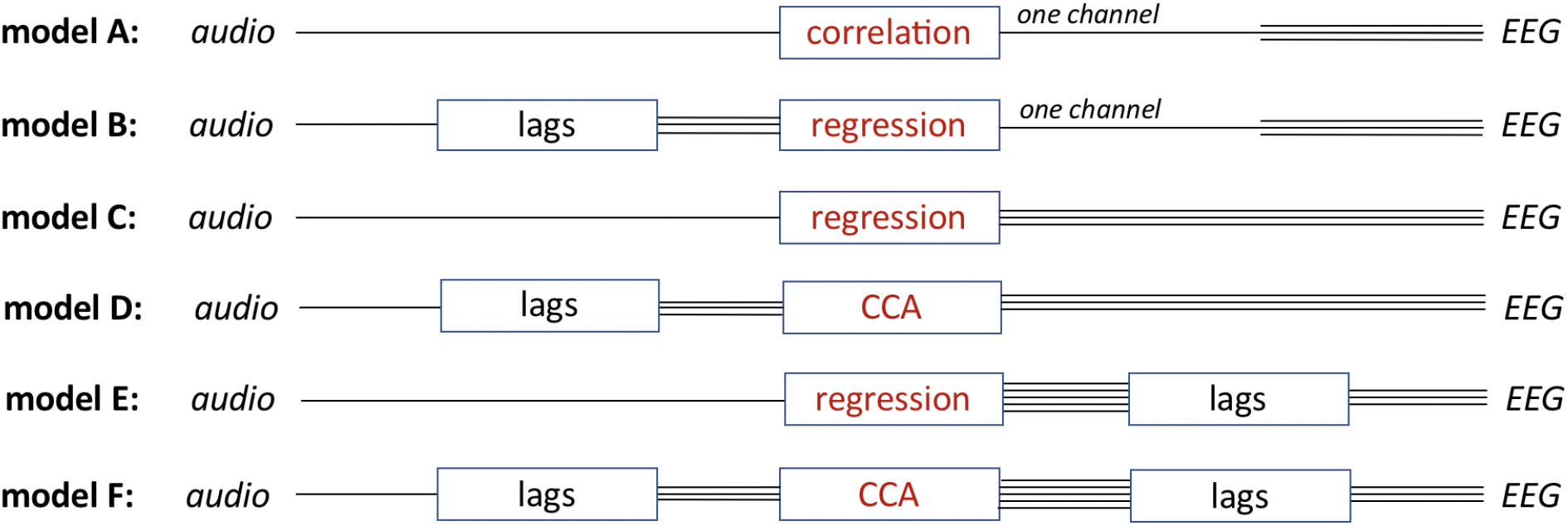
The six basic stimulus-response models considered in this study.

- *Model* **A** compares one EEG channel with the stimulus envelope, with no spatial or temporal filtering (*L*_*A*_ = 1, *L*_*X*_ = 1) other than the time shift *S* common to all models.
- *Model* **B** compares one EEG channel with a linear combination of time-lagged envelope signals (*L*_*A*_=11, *L*_*X*_=1) obtained by regression. This corresponds to a standard forward model as reported in the literature.
- *Model* **C** compares the envelope to a linear combination of EEG channels (without lags; *L*_*A*_ = 1, *L*_*X*_ = 1) obtained by regression. This is analogous to the basic backward model considered in de Cheveigné et al. (2018), or the single-delay model of Hausfeld et al. (2018)
- *Model* **D** compares linear combinations of time-lagged envelope signals with linear combinations of EEG channels (*L*_*A*_ = 11, *L*_*X*_ = 1), obtained by CCA. This is analogous to “CCA model 1” of de Cheveigné et al. (2018).
- *Model* **E** compares the envelope with a linear combination of time-lagged EEG channels (*L*_*A*_ = 1, *L*_*X*_ = 11) obtained by regression. This is analogous to the backward model of e.g. Fuglsang et al. (2017), or the multiple-delay model of Hausfeld et al. (2018).
- *Model* **F** compares linear combinations of time-lagged envelope signals with linear combinations of time-lagged EEG channels (*L*_*A*_ = 11, *L*_*X*_ = 11), obtained by CCA. This is analogous to “CCA model 2” in de Cheveigné et al. (2018).

To summarize the similarities and differences: models **A** and **B** relate the stimulus to just one of the *J* EEG channels. In contrast, all other models relate the stimulus to the ensemble of EEG channels. For models **A, B, C** and **E** the fit is based on univariate regression, and for models **D** and **F** on a multivariate CCA model. For univariate regression models, the fit is quantified by a single correlation coefficient, and for CCA by as many coefficients as CC pairs (Fig. 2).

Not counting *S*, the number of parameters in the fit is 1 for model **A**, *L*_*A*_ = 11 for model **B**, *J* = 64 for model **C**, *L*_*A*_ + *J* = 55 for model **D**, *JL*_*X*_ = 704 for model **E**, and *L*_*A*_ + *JL*_*X*_ = 715 for model **F**.

#### Model G

In addition to basic models **A**-**F**, we define a reference or “gold standard” model **G**, variant of model **F**, with a performance close to the best we found, and with a relatively straightforward and precisely defined implementation that can help future studies to document further improvements in performance. Details of this model are given in the Results section.

#### Display of results, statistics, implementation

Results are evaluated using the three metrics described above, and plotted as a function of selected parameters chosen to offer insight. Effects are tested for statistical significance using a non-parametric Wilcoxon signed rank test over subjects. Processing scripts in Matlab make use of the NoiseTools toolbox (http://audition.ens.fr/adc/NoiseTools/). Scripts are available at http://audition.ens.fr/adc/NoiseTools/src/NoiseTools/EXAMPLES/match_mismatch/.

## 2 Results

In the following, we evaluate and compare the models, focusing on the factors that affect performance. Section 2.1 compares performance of the six basic models (**A, B, C, D, E, F**; Fig 3), using the correlation metric for simplicity and to allow comparison with prior studies. Section 2.2 then introduces the MM classification task, and explores how sensitivity and error metrics depend on segment duration. Section 2.3 explores the dependency of all three metrics on the number of spatial dimensions (number of channels or principal components) and temporal dimensions (lags or filter channels). Based on this, Section 2.4 proposes model **G** for use as a comparison point in future studies. Section 2.5 investigates factors that cause the classifier to fail, and Sect. 2.6 summarizes performance across models.

**Figure 3:**
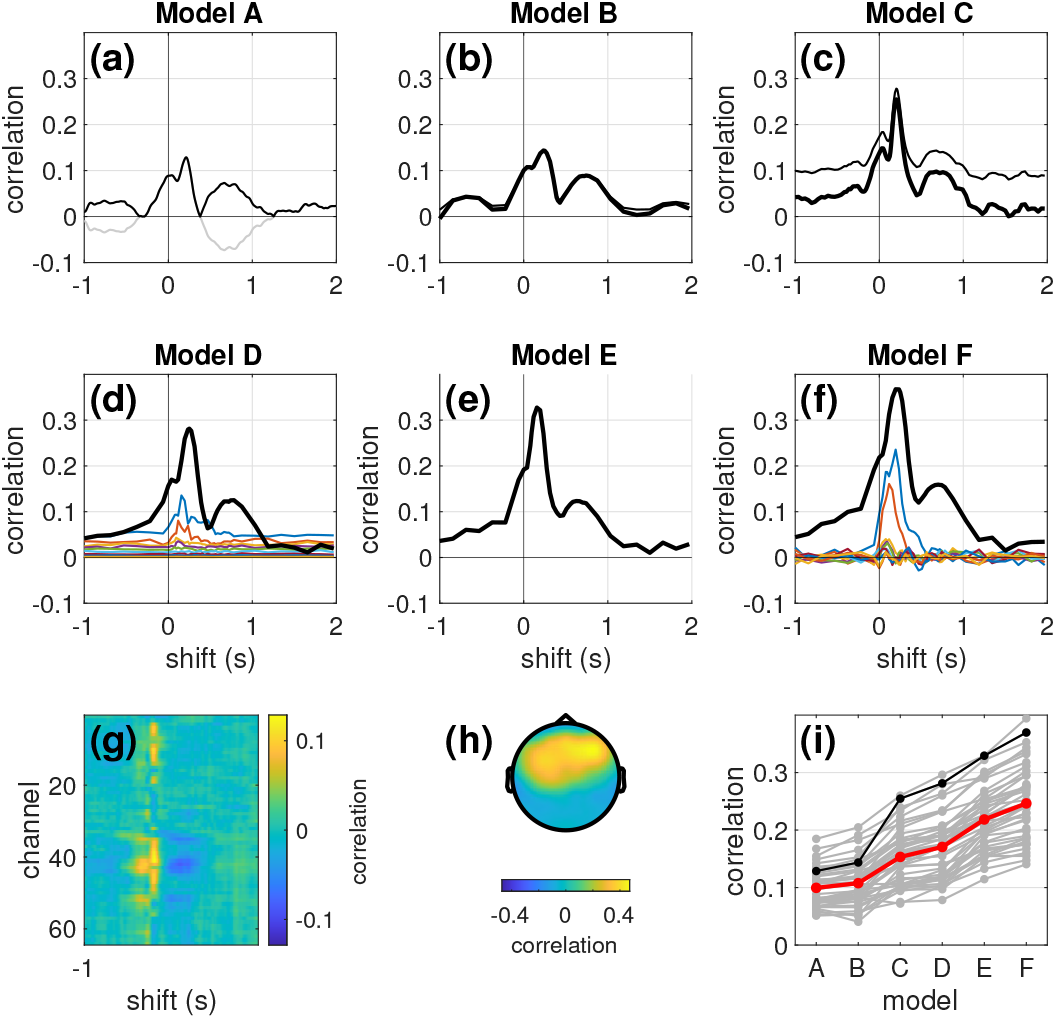
Baseline models **A** to **F** for one subject (subject 4, a relatively good subject selected for visual clarity). Panels (a)-(f): correlation as a function of overall time shift *S* for each model. (a) Cross-correlation function (gray), or absolute value of same (black) between channel 10 of EEG (FC3) and the stimulus envelope (Model **A**). (b) Correlation between channel 10 of EEG and the projection of channel 10 on the time-lagged stimulus, with crossvalidation (thick) or without (thin) (Model **B**). (c-f) Cross-validated correlation between stimulus-based and EEG-based components for Models **C**-**F**. (g) Cross-correlation function between the stimulus envelope and the EEG signal for each EEG channel (as in Model **A**). (h) Topography of correlation coefficients beween EEG-based component of first CC pair and individual EEG channels (model **D**). (i) Best correlation obtained for each model, for all subjects (gray). The average over subjects is in red, and the selected subject 4 is in black.

### 2.1 Correlation metric for basic Models

Figure 3 summarizes results obtained with the basic models. The first and second rows display correlation (calculated over the duration of each trial, ∼50 s) for models **A** to **F** for one subject (subject 4). Figure 3 (i) summarizes these results by plotting, for each model, the peak cross-validated correlation averaged over subjects (red) and for individual subjects (gray, black for subject 4).

#### Model A

This is the simplest incarnation of a stimulus-response model (Eq. 2), with **F** and **G** both identity transforms. Figure 3 (a) shows correlation (gray) between stimulus and EEG as a function of shift *S* for the best EEG channel (FC3). This is equivalent to the cross-correlation function between stimulus and response (positive and negative values). All other plots represent correlation of a signal *with its projection* (positive values only). To ease comparison between model **A** and the others, the plot also shows the absolute value of the cross-correlation function in black. The shape of the cross-correlation function differs slightly between electrodes (Fig. 3 (g)), implying that response properties are not uniform across the brain. Peak absolute correlation is 0.13 for this subject; peak values for other subjects can be read off Fig. 3 (i). For the best subject, a bit less than 4% of the variance of the best EEG channel is explained by the stimulus via this model.

#### Model B

The same EEG channel is projected onto the subspace spanned by the *L*_*A*_=11 time-lagged stimulus signals, yielding weights that define an optimized *FIR filter* applied to the stimulus. Figure 3 (b) shows correlation (thin) and cross-validated correlation (thick) as a function of shift *S* for the best channel (FC3). Cross-validated correlation differs only slightly from raw correlation (thick versus thin) suggesting minimal overfitting in this simple model. Peak correlation is greater than for model **A**, suggesting that the FIR filter has improved the fit. This improvement is robust across subjects (Fig. 3, (i)), as confirmed by a Wilcoxon signed rank test (p<10^*−*8^).

#### Model C

The stimulus is projected onto the subspace spanned by the *J* =64 EEG channels, yielding an optimized *spatial filter*. Peak cross-validated correlation between the stimulus signal and its projection (spatially-filtered EEG) is greater than for the previous two models across all subjects (Fig. 3 (c), p<10^*−*8^). The topography associated with the projection (correlation with individual EEG channels) shows a pattern typical of auditory responses (Fig. 3, (h)).

#### Model D

In this model, the time-lagged stimulus is related to the multichannel EEG using CCA. This results in multiple CC pairs, each quantified by a cross-validated correlation value (Fig. 3 (d)). The first CC is plotted in black, subsequent CCs are in color. Each CC is defined by a distinct FIR filter applied to the stimulus, and a distinct spatial filter applied to EEG. Multiple CCs suggest that the stimulus-response model captures multiple brain sources sensitive to different modulation frequency bands within the stimulus. Peak cross-validated correlation for the first CC is greater than for all previous models across subjects (p<10^*−*11^).

#### Model E

Time lags are applied to all EEG channels (but not the stimulus), resulting in a backward model in which the EEG undergoes both spatial and temporal filtering, analogous to the backward model of e.g. Fuglsang et al. (2017). Peak cross-validated correlation is greater than for all previous models across subjects (p<10^*−*6^).

#### Model F

Finally, lags are applied to both stimulus and EEG. Each CC then associates a distinct FIR filter applied to the stimulus with a distinct *multichannel FIR filter* applied to the EEG. Peak cross-validated correlation is again higher than all previous models across subjects (p<10^*−*12^).

Performance improves from model **A** to **F** for most subjects (Fig. 3 (i) gray lines). Three features seem to contribute to a better fit: *spatial filtering* leveraging the multichannel nature of EEG (models **C**-**F**), *temporal filtering* allowed by augmenting the data with time shifts (models **B**-**F**), and *CCA* which optimally relates multivariate representations of both stimulus and response (models **D** and **F**). It is worth noting that these models differ also in their number of free parameters, from 1 for model **A** (not counting shift *S*) to 735 for model **F** (see Methods). One might speculate that more parameters, rather than any particular feature, is what explains the progression in correlation scores. However, these results were obtained for cross-validated correlation for which overfitting should be detrimental. Thus, it seems that the more complex models genuinely provide a better fit, as confirmed with other metrics, below.

### 2.2 Classification-based metrics

We take the best model in terms of correlation (**F**), and rate its performance on the MM task in terms of metrics sensitivity and error rate, which are both based on the distribution of the difference *Δ*_*s*_ = *d*_*mm*_ − *d*_*m*_ *>* 0 between matched and mismatched segments of duration *D* (Methods).

Figure 4 (a) shows the distribution of *Δ*_*s*_ for *D*=10s (red) and *D*=1.25 s (blue). For longer segments, the distribution includes mostly positive values resulting in correct classification, whereas for shorter durations it includes a greater proportion of negative values. The degree to which the distribution is dominated by positive values, minimizing error, is captured by the sensitivity index (standardized mean of *Δ*_*s*_). Larger is better.

**Figure 4:**
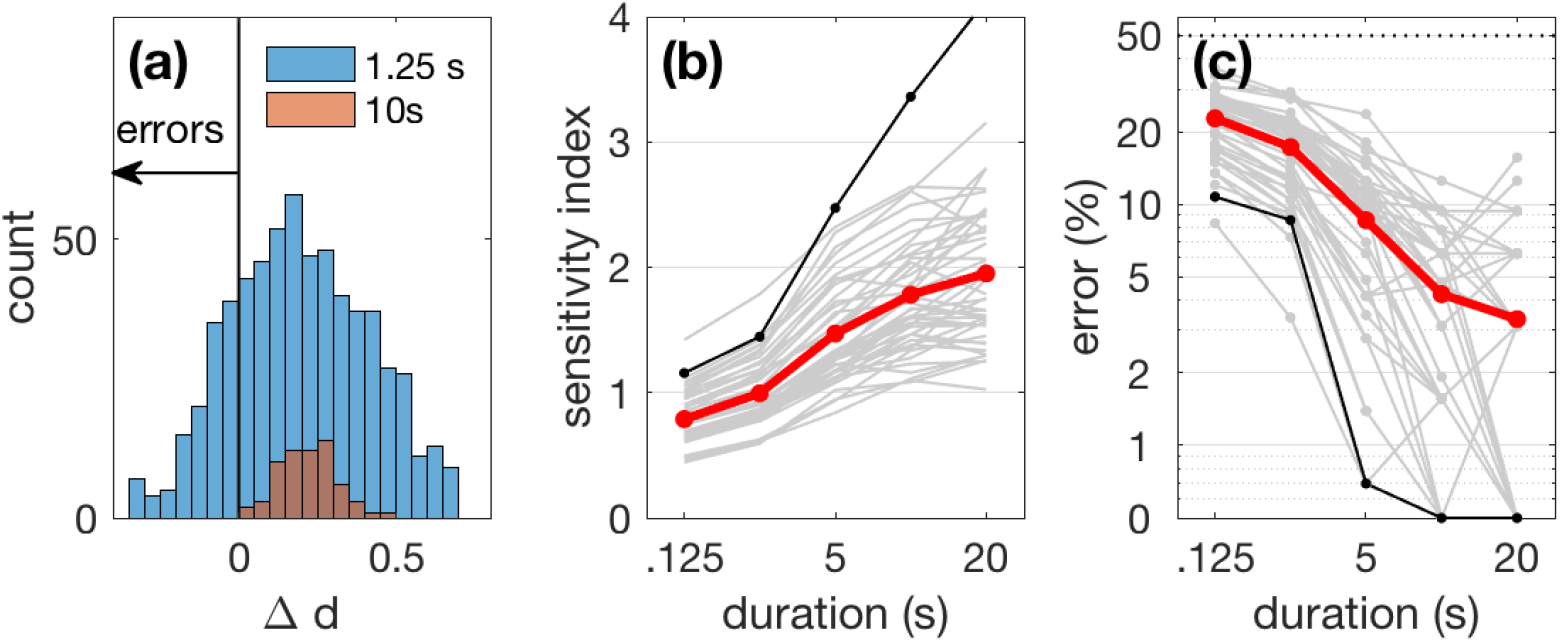
Match-mismatch task. (a) Histogram of Δ_*s*_ for segment durations of 1.25 s (blue) or 10 s (red), for subject 4. For shorter segments the histogram is wider and there are more errors. (b) Sensitivity index *µ/σ* as a function of segment duration averaged over subjects (red) and for each individual subject (gray, subject 4 is black), for model **F**. (c) Error rate.

Figure 4 (b) shows the sensitivity index as a function of segment duration averaged over subjects (red) and for individual subjects (gray, subject 4 is black). Figure 4 (c) likewise shows error rate as a function of duration. We expect the sensitivity index to be greater, and the error smaller, for a longer segment duration because the task is easier, and indeed this is the case. In the following we focus on *D*=5 s, for which the error rate averaged over subjects is ∼9% for this model (model **F**, *L*_*A*_ = *L*_*X*_ = 11). The variability over subjects is remarkable: at 5 s the error rate ranges from close to 0 (perfect classification) to more than 20%. Chance rate is 50%.

### 2.3 Spatial and temporal dimensionsionality

This section explores ways to further optimize performance. Comparing basic models (Fig. 3) it appears that performance can benefit from both spatial filtering and lags. However, a recurring issue for stimulus-response models is overfitting, which depends on the complexity of the model, function here of both the number of spatial dimensions (channels or PCs), and the number of lags. Both factors are explored here.

#### Number of spatial dimensions

Using model **C** (no lags) as a reference point, Fig. 5 (blue) shows the effect of applying PCA to the EEG data and discarding PCs beyond a certain rank *N*. The sensitivity index peaks, and the error rate is minimal, for *N* ≈ 32, suggesting that overfitting may be occurring due to excess dimensionality and that reducing dimensionality can mitigate its effects.

**Figure 5:**
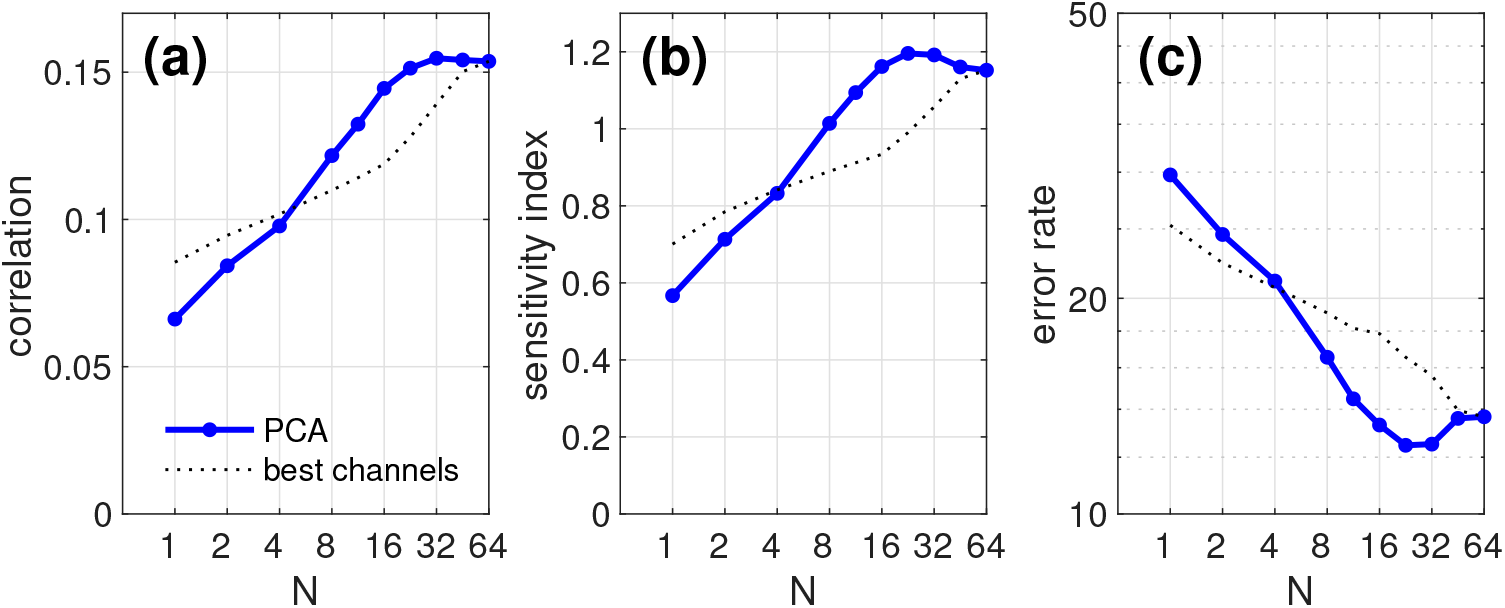
Performance as a function of the number of spatial dimensions, averaged over subjects. (a) Cross-validated correlation as a function of the number of PCs retained after PCA of the 64-channel EEG data. The dotted line represents subectaveraged correlation for subsets of EEG channels chosen for maximal correlation with the stimulus. (b) Sensitivity index. (c) Error rate. The model here includes no lags (similar to model **C**). Segment size is 5 s.

Truncating the series of PCs is markedly better than the simple expedient of discarding channels (dotted line; channels were sorted by decreasing correlation with the stimulus and the least correlated were discarded). This result is interesting in relation to claims that reducing the number of electrodes can yield equivalent performance to the full set, or even better performance due to less overfitting (Montoya-Martínez et al., 2021). Such is not the case here: the sensitivity index (Fig. 5 (b), dotted line) rises monotonically, implying that it is best to keep the full set. At no point does performance reach the level that can be attained by selecting PCs from a PCA applied to the full set of electrodes. The conclusion is simple: more electrodes *is* better.

#### Number of lags

Figure 6 shows metrics of correlation, sensitivity index, and error rate as a function of the number of lags (*L* = *L*_*A*_ = *L*_*X*_) averaged over subjects (red) and for individual subjects (gray, subject 4 is black). As the number of lags is increased, correlation and sensitivity increase until ∼ *L*=32 (250 ms), then decrease beyond. This peak is mirrored by a dip in error rate at *L* = 32. The best error rate is 2.8% on average over subjects.The reversal beyond *L* = 32 might reflect an increase in *d*_*mm*_ relative to *d*_*m*_, thus reducing the numerator *m* in Eq. 3, or an increase in their variablity, thus increasing the denominator *σ*.

**Figure 6:**
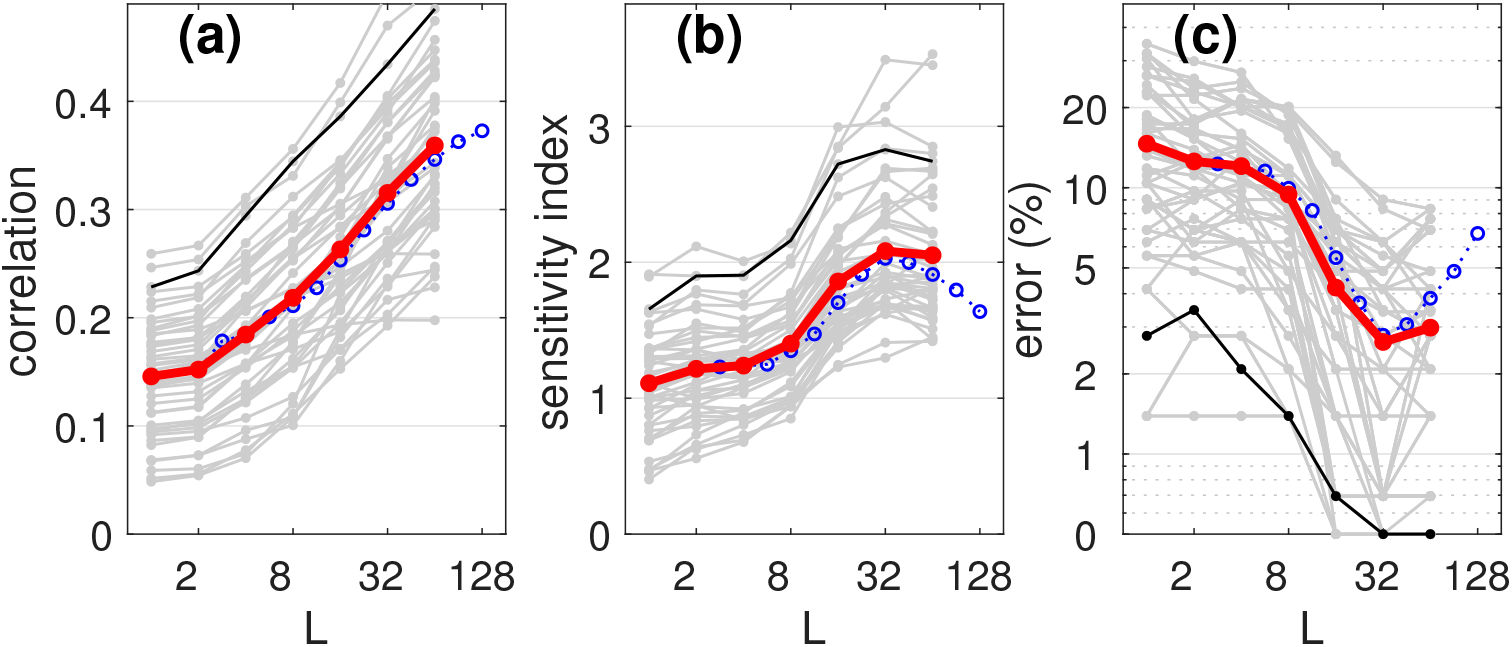
Performance as a function of the number of lags (or the order of the dyadic filters) applied to both stimulus and EEG (*L*=*L*_*A*_=*L*_*X*_). (a) Crossvalidated correlation as a function of *L* averaged over subjects (red) and for all subjects (gray, black is subject 4). Blue symbols are for a dyadic filter bank instead of lags (see text). (b) Sensitivity index. (c) Error rate. Segment size is 5 s.

A fall in performance with larger *L* can be a sign of overfitting, as a model with many parameters can fit the minutiae of the training data but generalize poorly to the test data However, the drop might also have a different cause, for example long lags might capture slow patterns that do not generalize well. We can arbitrate between these explanations by comparing results for lags 1 *… L* (Fig. 6, red) with those for a dyadic filter bank with FIR filters of order *L* (blue). A filterbank of *L*^*′*^ dyadic filters of order *L* has fewer channels (*L*^*′*^<*L*) and thus requires fewer parameters in the model (*L*^*′*^ = 10 for *L* = 32; *L*^*′*^ = 12 for *L* = 64, etc.). Thus, the drop should occur for larger *L* for the dyadic filterbank (blue) than for lags (red) if it reflected the number of parameters. Instead it occurs at the same value of *L*, suggesting indeed that performance drops because longer lags capture slow patterns that do not generalize well.

In a previous study (de Cheveigné et al., 2018) we obtained better performance with the dyadic filterbank than with lags. We do not replicate that result here: performance is similar for both for equal *L*. Once again, the variability of these metrics over subjects is remarkable. For *L*=32, the error rate for 5-second segments ranges from 0% for the best 10 subjects to ≈9 % for the worst. Incidentally, the error rate averaged over hearing-impaired subjects (1.7%) is smaller than for normal hearing subjects (3.4%), p<0.001, t-test, as was found in other studies (Goossens et al., 2018; Decruy et al., 2020; Fuglsang et al., 2020).

### 2.4 Model G (“gold standard”)

Based on these results, we define more precisely one particular model, using one set of parameters that produces close to state-of-the-art performance on a publicly available dataset, to serve as an easy-to-implement reference with which to evaluate new algorithms. This reference model and evaluation are described in detail to allow replication.

Model **G** involves the following steps: (1) a time shift *S* of 200 ms is applied to the EEG to advance it relative to the audio envelope, (2) data are preprocessed as described in Methods, (3) PCA is applied to the EEG and the first 32 PCs are selected, (4) audio envelope and EEG are both augmented by applying lags 0*…* 31 (i.e. a 32/128=250 ms range of lags), (5) the augmented data are fit by a linear stimulus/response model based on CCA, (6) sensitivity and error rate metrics estimated using segments of duration 5 s.

To be precise: for each crossvalidation fold, the CCA solution is trained on a subset of 15 trials and tested on the 16th (left out). All consecutive 5 s segments of audio within the left-out trial are considered, and for each the Euclidean distance *d*_*m*_ between that segment of audio and the corresponding segment of EEG is calculated (matched distance), as well as the average Euclidean distance *d*_*mm*_ between that audio segment and all EEG segments of all 15 other trials (mismatched distance), yielding a difference score Δ_*s*_ = *d*_*mm*_ − *d*_*m*_ for that segment. Values of Δ_*s*_ are aggregated over segments and folds. The ratio between the mean of the distribution of Δ_*s*_ and its standard deviation yields the *sensitivity* metric. The proportion of samples for which Δ_*s*_ falls below 0 yields the *error rate* metric. Distance calculations take into account the first 5 CCs of the CCA solution with equal weights^1^.

To evaluate a new method, the recommended procedure is (1) install model **G** on the same system as the new algorithm, (2) test it using the same publicly available database as we use, and verify that the metrics yield scores consistent with what we report, and (3) apply the new method to that database and compare scores with (2). The reason for step (2) is to control for installation-specific differences (e.g. single versus double precision, etc.).

Alternatively, if a different database is to be used, do (1) as above, then (2’) test model **G** using the new database, and (3’) test the new method on the new database and compare scores with (2’). In any event, it is not recommended to compare a new method with prior methods on a different database, or with different metrics, or with a different task. Thus, there would be little merit in comparing the scores we report here to those reported in the literature for AAD.

### 2.5 Anatomy of an error

One of our goals is to gain a better understanding of factors that determine model performance. An error occurs when the difference Δ_*s*_ = *d*_*mm*_ − *d*_*m*_ falls below zero, and this might be caused by a relatively small value of *d*_*mm*_ or a relatively large value of *d*_*m*_. It is clear from Fig. 7 that the latter is the main factor for this subject (subject 3, relatively poor model performance). The top panel shows *d*_*m*_ (dots) and *d*_*mm*_ (crosses) for all segments of all trials. The mismatched distances are distributed tightly around *d*_*mm*_ ≈ 1.4 as expected (Sect. 1.1, Metrics) whereas matched distances *d*_*m*_ are mostly smaller, as clear also from the scatterplot of *d*_*m*_*m* versus *d*_*m*_. The diagonal line in Fig. 7 (b) represents the classification boundary Δ_*s*_ = 0: all points to the right and below this line (red) are misclassified.

**Figure 7:**
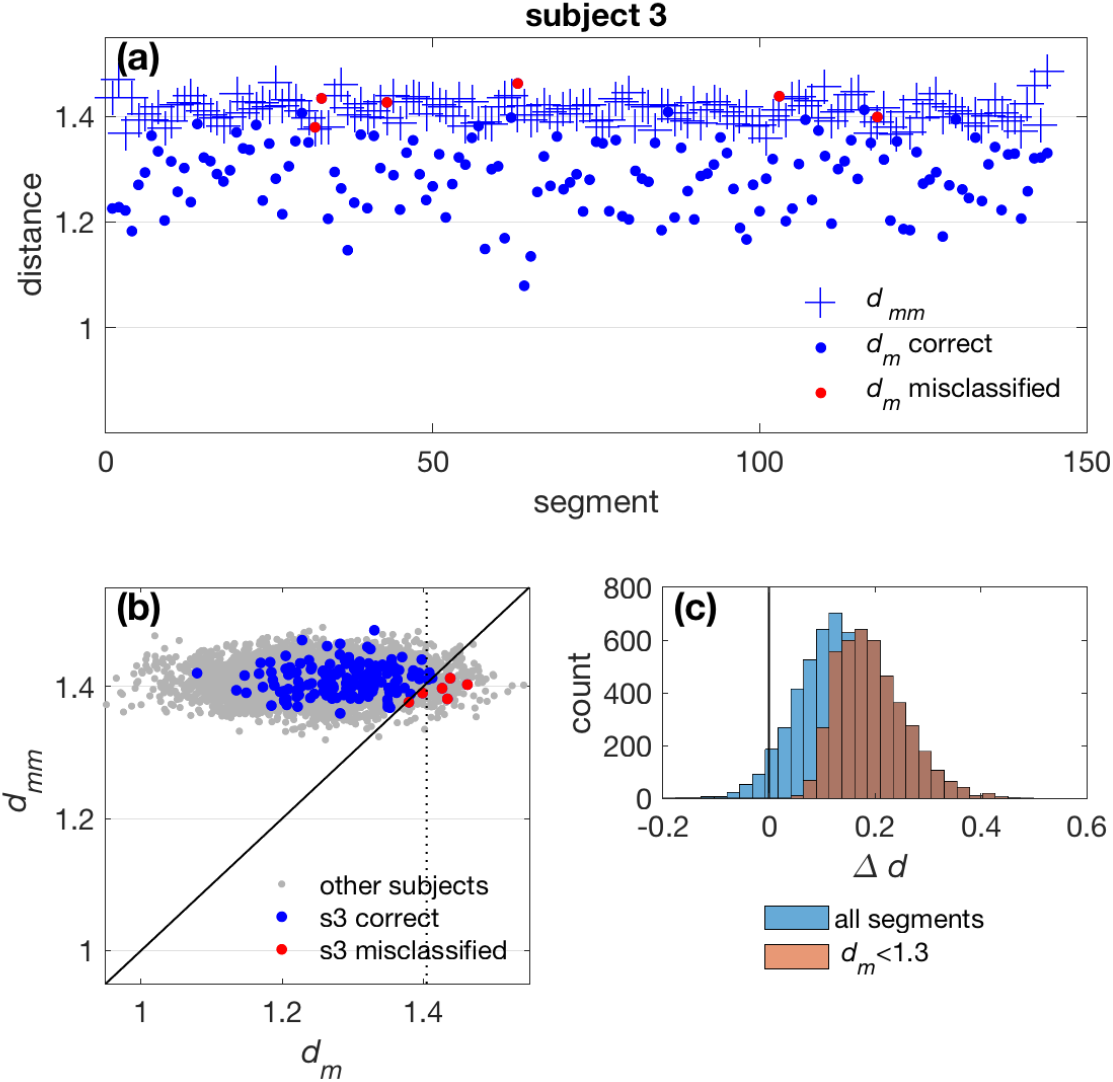
(a) Euclidean distance between matched (dots) and mismatched (+) segments of duration 5 s for all trials of one subject (subject 3, chosen for a relatively high error rate). Red dots indicate classification errors. (b) Scatterplot of mismatched versus matched distances for subject 3 (blue/red) and all other subjects (gray). The diagonal represents the classification boundary Δ_*s*_ = 0. Points below that line (red) are misclassified. (c) Histograms of values of Δ_*s*_ for all segments (blue), and for segments for which the matched distance *d*_*m*_ is less than 1.3 (brown).

The matched distance *d*_*m*_ is a good predictor of classification reliablity: for *d*_*m*_ *<* 1.3 the classification statistic Δ_*s*_ is distributed far from the decision boundary (Fig. 7, (c) brown). Classification is highly reliable in that case, whereas for larger values of *d*_*m*_ the classification is less reliable. This implies an asymmetry in the conclusions that can be drawn from the classifier. For example a hypothetical “attention-monitoring” device might rapidly and reliably detect that a stimulus *has* registered within a subject’s brain, but the opposite conclusion that it *has not* registered would take longer and/or be less reliable. Reliability is a useful adjunct to the decision.

What factors might inflate *d*_*m*_? Regression on the RMS of the EEG signal shows a significant but weak correlation with response power (r=0.12, p<10^*−*7^), suggesting that high-amplitude glitches in the EEG might be a contributory factor. On the other hand, a significant but weak negative correlation with RMS stimulus (r=-0.07, p<10^*−*20^) suggests a possible small contribution of lulls in the stimulus. However, these small correlation values suggest that other factors, unknown, dominate.

### 2.6 Summary of methods

Figure 8 summarizes error rates obtained with each of the models **A**-**G**, averaged over subjects. Models **A** and **B** are classic forward models that attempt to predict one channel of EEG from the stimulus representation. Models **C** and **E** are classic backward models that attempt to infer the stimulus representation from the EEG. Models **D, F** and **G** are hybrid models based on CCA. The best model (**G**) makes an order of magnitude fewer mistakes than the worst (**A**). For a 5 s window the error rate for model **G** is less than 3% on average over subjects (0% for 10 subjects). Extrapolating from progress so far, we think that further progress is possible. Associated with the publicly available dataset that we used, model **G** might serve as a “gold standard” for comparative evaluation of such future progress.

**Figure 8:**
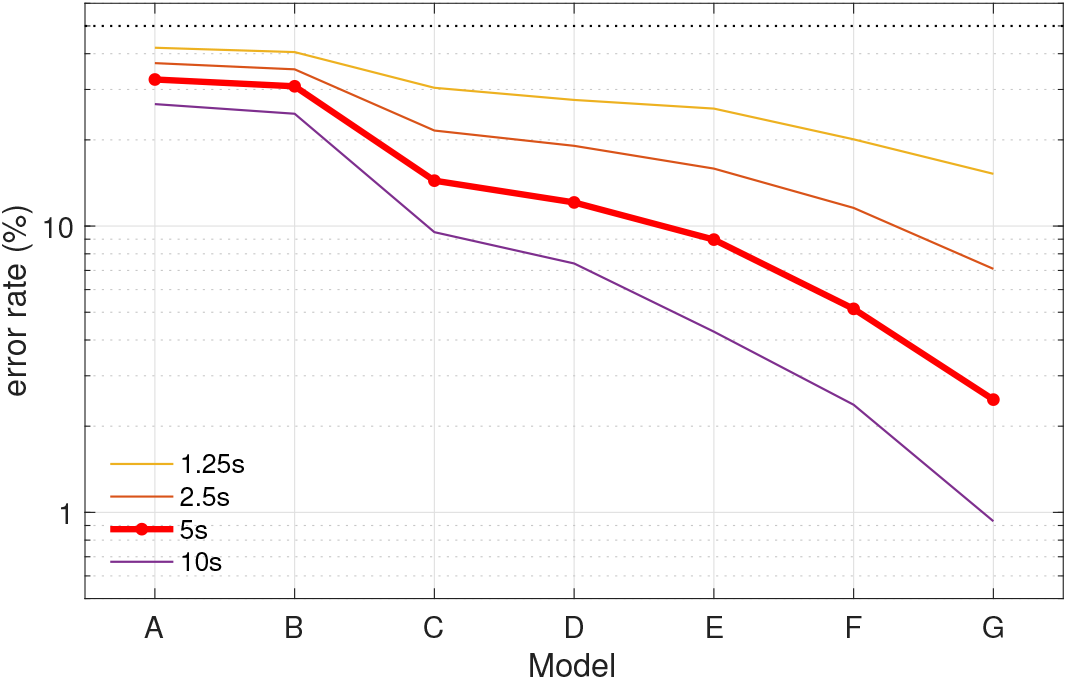
Summary of error rates for models **A**-**G**, averaged over subjects, for several values of duration *D*. The dotted line represents chance (50%). At each duration, model **G** makes fewer errors than its closest competitor (**F**, p*<*10^*−*4^, Wilcoxon signed rank test).

## 3 Discussion

This study offers two main contributions. First, it introduces a simple objective task to help in the evaluation of stimulus-response models. Second, it documents a set of techniques that boost performance beyond state of the art.

### The need for an objective task

A straightforward quality metric for a stimulus-response model is *correlation*, between response and predicted response in a forward model, between stimulus and inferred stimulus in a backward model, or between transforms of both in a hybrid model. That metric is simple and informative: better models tend to yield higher scores. However an elevated score can also result from chance correlations. These are more widely distributed for data dominated by low frequencies, which could mislead a researcher to conclude that lowpass filtering, for example, improved the model despite the loss of relevant information carried by higher frequencies (Kriegeskorte and Douglas, 2019). An objective task alleviates this problem, because loss of relevant information must impair task performance. Another argument in favor of an objective task is that it is a better measure of the model’s “real world” value.

Why three metrics? Firstly, they are not equivalent: referring to Fig. 6, correlation increases monotonically with lag (a), whereas sensitivity and error rate show a reversal at *L* = 32 (b, c). *Error rate* is directly relevant for applications but somewhat coarse and brittle (it depends on a few samples near the classification boundary). *Sensitivity* depends on all samples by summarizing them based on their mean and standard deviation, but like error rate it requires a task. The appeal of *correlation* is that it is task-agnostic. Thus, the three metrics are complementary.

### Selective versus sustained attention

Auditory attention is often investigated in a situation where multiple stimuli compete for attention, for example concurrent pulse trains (Hillyard et al., 1973), or competing voices (Kerlin et al., 2010), or competing instruments (Treder et al., 2014). It may also be probed as a difference in response to a stimulus in the presence, or absence, of concurrent visual stimulation (Molloy et al., 2015), or of a behavioral task (Scheer et al., 2018). In each case, a comparison is made between brain responses to the same stimulus recorded in two situations. In contrast, the MM task requires only a single recording, and, more importantly, assumes no competition for auditory resources. As such it might be of use to monitor the general attentive (versus inattentive) state of a subject, for example to determine whether an alert has been perceived, or a message has registered, or to detect drowsiness, or evaluate the minimal conscious state in patients in locked-in state, or to detect if a participant is actively paying attention to continue/pause a BCI application.

The AAD task is relevant for BCI applications such as cognitive control of acoustic processing in a hearing aid. However, even for those applications it may be fruitful to optimize the stimulus-response models using the MM task. We speculate that improvements obtained for the simpler MM task may transfer to the harder AAD task. A practical issue with AAD is that it relies on specific experimental setups with competing voices, attention task instructions, and greater demands for listening effort. The MM task does not rely on predefined data labels but instead derives them (match versus mismatch) from a manipulation of the input data. It can therefore be applied to any dataset of brain responses to sound. An analogous approach has been used successfully for self-supervised learning, for instance, by training neural networks to predict whether video and audio segments are temporally aligned (Owens and Efros, 2018; Arandjelovic’ and Zisserman, 2018). The task and metrics are applicable to self-supervised training of large-scale neural networks that require extensive training sets. Being free of reliance on particular ‘attention labels’, the MM-approach is better suited to evaluate and compare models across datasets with different experimental setups.

Another practical issue with AAD is potential mislabeling due to attentional capture by the wrong stream. We cannot be sure that a subject consistently followed the instructions throughout, and thus a certain proportion of the database might be *mislabeled*, an important concern when evaluating well-performing models. The MM task is thus appealing as it can be evaluated on data with only one sound stream. A downside is that MM is blind to potential brain processes specific to attention that AAD might capture. The two tasks are thus complementary.

### Encoding, decoding, and hybrid models

A forward (encoding) model is judged by the proportion of brain signal variance that it can account for (Naselaris et al., 2011; Kriegeskorte and Douglas, 2019). However, much of brain activity is not stimulus-related, so that proportion is small even for a model that perfectly predicted all stimulus-related brain activity. Analogous comments can be made with respect to backward models (stimulus reconstruction) that infer only select aspects of the stimulus representation rather than its entirety.

The appeal of hybrid models such as CCA (Dmochowski et al., 2017; de Cheveigné et al., 2018; Zhuang et al., 2020) is that both stimulus and EEG are stripped of irrelevant variance, leaving remainders that can more usefully related one to the other. The model then is predictive of a *transform* of the measured brain response, rather than of the response itself, which makes it harder to interpret than a forward model. For example, Model **F** defines a set of linear transforms of the time-lagged EEG signals, each predicted from the stimulus envelope via an FIR filter, which harder to interpret than Model **B** that directly predicts an EEG channel, or even Model **D** that predicts spatially filtered EEG.

The upside of hybrid models is that the transformed response **XG** (right hand side of Eq. 2) arguably offers a closer (less noisy) view of the information coded by sensory-dependent parts of brain activity (Kriegeskorte and Douglas, 2019). Equation 2 can be generalized to more complex transforms *f* (*A*) and *g*(*X*) (e.g. Andrew et al., 2013), and it may be useful to allow *f* to depend on *X* (e.g. sensory processing dependent on brain state) and *g* to depend on *A* (e.g. a different model of brain activity in response to speech and music).

### What makes a good model?

Prior studies using a forward model (similar to model **B**) or a backward model (similar to **C** or **E**) typically report performance that is “above chance” but still rather poor. For example, a score of r=0.1 to 0.2 means only 1 to 4% of the variance is explained, and furthermore a correct-classification score of 90% for a segment of 60 s duration (typical subject of O’Sullivan et al., 2015), implies a decision delayed by one minute and wrong on one trial out of every ten. For applications, it is crucial to achieve shorter latency and better reliability, and from the scientific perspective it is desirable to find models that allow a better fit to the data.

CCA allows both data streams to be stripped of irrelevant variance, resulting in a better fit as reflected by higher values of the correlation metric (compare models **C** versus **D**, or **E** versus **F**) and the multiple correlation coefficients support multivariate classification, with a further boost to task-based metrics.

An important ingredient in the more successful models is *lags*, that allow the algorithms to synthesize FIR or multichannel FIR filters that can absorb convolutional mismatch between the stimulus and response, thus resulting in better performance (compare models **A** versus **B, C** versus **D**, or **E** versus **F**). Adding lags increases the dimensionality of the data space, which is beneficial *as long as the optimal transforms can be found*. If not, due to overfitting, the larger dimensionality may instead by harmful.

Model overfitting was addressed here using *dimensionality reduction* which can be achieved trivially by discarding sensor channels (with limited success, c.f. dotted line in Fig. 5), or limiting the number of lags (with greater success, Fig. 6 center and right). Replacing the set of lags by a smaller number of channels of a dyadic filter bank also reduces dimensionality, with a considerable reduction in computation cost but little difference in performance (compare red and blue lines in Fig. 6). Applying PCA and selecting a subset of components also reduces dimensionality, with a slight boost in performance (Fig. 5) (see also de Cheveigné, 2021). An additional benefit is to reduce computational cost, which can otherwise become prohibitive if many lags are introduced (the bottleneck is an eigendecom-position which costs *o*(*N* ^3^)).

The reduction in performance beyond *L* = 32 (∼250 ms) that we observed for this dataset (Fig.6) suggested overfitting, which could merely result from a larger number of free parameters, or from the fact that higher-order FIR filters can enhance slow patterns (low frequencies) that don’t generalize from training data to test. The latter seems more likely since replacing *L* lags by a smaller number of dyadic filters of order *L* had little impact on performance (compare blue to red in Fig. 6). The knee occurs at the same value of *L* (32), suggesting that filter order (or lag span) is the critical factor.

With a ∼200 ms shift and a maximum lag of ∼250 ms, the model associates stimulus samples with response samples that occur up to ∼450 ms later. However, we cannot use this to make a strong statement concerning brain processing latencies because of the potential smearing effect of the filters applied in preprocessing (Sect. 1.3) (de Cheveigné and Nelken, 2019).

#### Whither now?

Further boosts in performance are needed to enhance the feasibility of potential applications. Based on what we know so far, there are several directions worth pursuing.

One is to improve the stimulus representation. Here, we used a rather crude representation, the stimulus envelope. Richer representations such as auditory filterbank (Biesmans et al., 2017), higher-order linguistic structure (Di Liberto et al., 2015), onsets (Oganian and Chang, 2019), or voice pitch (Forte et al., 2017; Teoh et al., 2019), etc. have been explored but remain to be developed further and integrated. Multi-set CCA (MCCA), which allows merging EEG across subjects, may ease development of such stimulus representations (de Cheveigné et al., 2019).

A second direction is to extract more information from the brain response. Typical models (including those reported here) exploit low-frequency components, but useful information may also be carried by high-frequency power (Synigal et al., 2020; Forte et al., 2017; Teoh et al., 2019). Standard linear techniques (such as CCA) are not directly applicable to enhance weak sources of power, but it may be possible to use quadratic component analysis (QCA) for that purpose (de Cheveigné, 2012). This entails forming cross-products between channels and/or lags, leading to very high-dimensional data for which an appropriate dimensionality-reduction strategy is crucial.

A third direction is better integration of information over time. As Fig. 7 (top) shows, errors occur only for segments for which the mismatch *d*_*m*_ is large, and these occupy only a small fraction of the time axis. A better understanding of what triggers large-mismatch events might allow them to be mitigated. Alternatively, since they are flagged by a high value of *d*_*m*_, it may be possible to integrate over a high-reliability (low *d*_*m*_) context to offer the application a reliable decision.

A fourth direction is more prosaic: better preprocessing, filtering, artifact rejection, parameter tuning, etc. Performance metrics are sensitive to preprocessing parameters, but no attempt was made to tune them in this study.

Finally, a fifth direction is to use more recent machine-learning methods in lieu of expertise-based approaches, in the faith that they will discover the same regularities and structure as embodied by hand-crafted methods, and more. Results so far are modest (Ciccarelli et al., 2019; Jalilpour Monesi et al., 2020; Tian and Ma, 2020; Das et al., 2020), but success in other fields suggests that machine-learning approaches are well worth pursuing.

## Acknowledgements

This work was supported by grants ANR-10-LABX-0087 IEC, ANR-10-IDEX-0001-02 PSL, and ANR-17-EURE-0017. Jens Hjortkjaer and Søren A. Fuglsang were supported by the Novo Nordisk Foundation synergy Grant NNF17OC0027872 (UHeal). We appreciate many helpful discussions with Jonathan Berent and his team. We gratefully acknowledge useful comments from two reviewers on earlier versions of the manuscript.

## Footnotes

1 A Matlab implementation is available at http://audition.ens.fr/adc/NoiseTools/src/NoiseTools/EXAMPLES/match-mismatch/

